# Transcranial alternating current stimulation at 10 Hz modulates response bias in the Somatic Signal Detection Task

**DOI:** 10.1101/330134

**Authors:** Matt Craddock, Ekaterini Klepousniotou, Wael el-Deredy, Ellen Poliakoff, Donna Lloyd

**Author notes:** Correspondence concerning this article should be addressed to Donna Lloyd, School of Psychology, University of Leeds, Leeds, LS2 9JT, UK.

## Abstract

**Background:** Ongoing, pre-stimulus oscillatory activity in the 8-13 Hz alpha range has been shown to correlate with both true and false reports of peri-threshold somatosensory stimuli. However, to directly test the role of such oscillatory activity in behaviour, it is necessary to manipulate it. Transcranial alternating current stimulation (tACS) offers a method of directly manipulating oscillatory brain activity using a sinusoidal current passed to the scalp.

**Objective:** We tested whether alpha tACS would change somatosensory sensitivity or response bias in a signal detection task in order to test whether alpha oscillations have a causal role in behaviour.

**Methods:** Active 10 Hz tACS or sham stimulation was applied using electrodes placed bilaterally at positions CP3 and CP4 of the 10-20 electrode placement system. Participants performed the Somatic Signal Detection Task (SSDT), in which they must detect brief somatosensory targets delivered at their detection threshold. These targets are sometimes accompanied by a light flash, which could also occur alone.

**Results:** Active tACS did not modulate sensitivity to targets but did modulate response criterion. Specifically, we found that active stimulation generally increased touch reporting rates, but particularly increased responding on light trials. Stimulation did not interact with the presence of touch, and thus increased both hits and false alarms.

**Conclusions:** tACS stimulation increased reports of touch in a manner consistent with our observational reports, changing response bias, and consistent with a role for alpha activity in somatosensory detection.

---

There is a wide range of evidence across multiple sensory modalities that spontaneous, ongoing neural oscillations in the alpha band - 8-13 Hz - have a direct role in perception and determining which stimuli are detected and which missed [1–5]. Much of this evidence is necessarily correlative, based on observations recorded using magneto- or electroencephalography (M/EEG). More direct evidence of causation requires direct manipulation of the ongoing oscillatory rhythms naturally and spontaneously exhibited by the brain.

Transcranial electrical stimulation (tES) offers one such method of directly influencing ongoing brain activity [6]. Three commonly used tES methods are transcranial direct current stimulation (tDCS), transcranial alternating current stimulation [7], and transcranial random noise stimulation (tRNS). Of these, tACS is particularly promising as a method by which to interact with endogenous rhythms, since it allows application of a sinusoidal current at a desired frequency. Indeed, there are several reports that tACS stimulation at or around 10 Hz modulates alpha power, increasing it even after stimulation has ended [8–10]. Furthermore, modulation of alpha oscillations using tACS also influences detection of visual targets phasically [8], consistent with the pattern found previously in the absence of tACS stimulation [11–13].

Effects of tACS on other sensory modalities, including audition [5] and pain [14], have been reported. Most relevant here, however, is how tACS stimulation may influence somatosensation. As in vision, tactile detection can be vary with the power of alpha oscillations recorded over somatosensory regions [3]. We found that detection of peri-threshold tactile stimuli was predicted from alpha power in a period shortly before stimulus onset [3]. In that study, participants performed the Somatic Signal Detection Task [15], in which they were asked to detect brief somatosensory stimuli delivered to their left index finger at detection threshold. Brain activity was simultaneously recorded using EEG. We found that power in the alpha frequency band influenced both true and false reports of somatosensory perception. As pre-stimulus alpha power increased, the probability of reporting touch decreased, both in the presence and absence of target stimuli. Given that alpha plays a similar role in both visual and tactile detection, and that alpha tACS modulates visual detection, it follows that manipulation of alpha using tACS may also modulate somatosensory detection.

A study by Gundlach, Müller, Nierhaus, Villringer, and Sem [16] found evidence consistent with this suggestion. They had participants perform a somatosensory detection task before, during, and after active alpha or sham tACS stimulation delivered over bilateral somatosensory cortices. Tactile stimuli were delivered to the participants’ right index finger. The intensity of the stimuli was continuously varied, but maintained at detection threshold using a staircase procedure. Detection thresholds for the stimuli in the periods before, during, and after the stimulation period did not differ on average. However, during active stimulation, detection thresholds varied in a phasic manner. Detection thresholds at opposite phases of the driving oscillations differed from baseline (pre-stimulation) performance in opposing fashion: some phases were associated with decreased thresholds while others were associated with increased thresholds.

However, a limitation of Gundlach et al.’s study [16] was that stimuli were always present. Thus, it is impossible to determine whether the changes in detection performance they observed were related to genuine variation in tactile sensitivity. TACS stimulation in the alpha frequency range may also induce faint tactile sensations contralateral to the stimulated region [17], which might increase false reports of touch during stimulation. A typical way of assessing performance on detection tasks is to calculate signal detection measures [18], which account for both hit rates - correct detection of target stimuli - and false alarm rates - false reports of target stimuli when the stimulus is absent. Sensitivity (*d′*) describes the ability to discriminate signal from noise. Response criterion (*c*) describes the degree of bias towards responding that a signal is present or absent.

In signal detection terms, the pattern of results reported in Craddock et al.[3] is consistent with changes in response criterion rather than sensitivity, since alpha power shifted hit and false alarm rates in the same direction. In addition, Gundlach, Müller, Nierhaus, Villringer and Sehm [19] reported that the somatosensory alpha rhythm decreased in power after tACS stimulation. Thus, in accordance with our results, decreases in power should increase reporting rates for touch, increasing both false alarms and hit rates, and thus not increase somatosensory sensitivity per se [3]. TACS stimulation might then change response criterion, biasing participants towards or against reporting stimuli, rather than changing sensitivity or detection threshold. Therefore, in order to test whether alpha tACS stimulation would induce changes in response bias, we had participants perform the SSDT while undergoing tACS.

## Material and methods

### Participants

Twenty-one right-handed participants (19 female, two male; ages: *µ* = 19.7 years, *σ* = .097) were recruited from the undergraduate population of the University of Leeds. Five additional participants were excluded following initial screenings for contraindications to receiving tACS stimulation (e.g. unremovable facial piercings, history of migraines). Participants received course credit or cash vouchers for participation. The study was approved by the ethical committee of the School of Psychology at the University of Leeds (ethics reference: 16-0019). All participants reported normal or corrected-to-normal vision and no tactile sensory deficits, and gave fully informed written consent.

### Apparatus

The stimulus array comprised a soft foam block in which a piezoelectric tactile stimulator (PTS) was embedded (Dancer Design, St. Helens, UK), with a red light-emitting diode (LED) attached next to the PTS. Participants placed their left index finger on top of the PTS. Tactile stimuli were produced by an auditory signal delivered from the experimental PC to the tactile amplifier (TactAmp 4.2, Dancer Design). Note that vibrations from the PTS were entirely inaudible when embedded in the foam block. A monitor located behind the stimulus array delivered instructions and visual cues. Participants sat approximately 70 cm in front of the monitor, with the stimulus array to the left of their midline. Participants responded with a button box held in their right hand. Timing and presentation of the stimuli was controlled using EPrime 2.0.

#### Transcranial alternating current stimulation (tACS)

Transcranial alternating current stimulation was applied using a neuroConn DC-Stimulator-Plus (Eldith, Neuroconn, Ilmenau, Germany). Two rubber electrodes (5cm by 5cm) in foam sponges - pre-soaked in saline solution - were placed over positions CP3 and CP4 of the international 10-20 electrode placement system. The sponges and electrodes were held in place using a rubber strap.

### Procedure

All participants took part in two experimental sessions separated by at least two days. Before beginning the experiment, the tACS montage was set up as above. The experiment itself was split into two parts. In the first part, each participant’s sensory threshold (i.e., 50% detection rate) was established using a two-alternative forced choice adaptive staircase procedure. Participants were given a series of trials consisting of two consecutive 1420 ms time periods. Each time period began with a green arrow presented for 400ms on the left side of the monitor and pointing down towards the participant’s finger. The numbers “1” and “2” were written on arrows marking the start of the first and second periods respectively. After the offset of each arrow, the screen remained blank for 1020 ms. On each trial, a 20 ms tactile pulse was delivered 500 ms after the offset of either the first or second arrow. After both time periods had elapsed, participants were prompted on screen to press button 1 or 2 on the button box to report whether the stimulus had been presented in the first or second time period. A further 1000 ms elapsed before the start of a new trial. Trials were repeated until a stable 50% detection threshold was reached or up to a maximum of 150 repetitions (no participant exceeded this maximum). Participants did not receive feedback.

In the main experiment, participants were asked to detect brief 20 ms tactile pulses delivered at sensory threshold. In the sham condition, random noise stimulation was applied for 30 s at 1.5 milliamps (mA). In the active condition a 10 Hz alternating current was delivered at 1.5 mA for 25 minutes (the approximate length of the experiment). The order of stimulation conditions was counterbalanced across subjects. In both conditions, stimulation ramped up from zero to 1.5 mA over 30 s at the beginning, and sloped back down to zero over 10 seconds at the end. At the start of each trial, a green arrow pointing down towards the participant’s left index finger appeared for 500 ms. This was replaced with a blank screen for 1 to 1.5 s. This was followed by a 20 ms stimulus period. In this period, there were four possibilities. On half the trials, a touch was delivered using the PTS. On half of those trials, the red LED flashed simultaneous with the occurrence of the tactile stimulus. On the remaining half of the trials, no touch was delivered, but the red LED flashed on its own on half of those trials. There were 204 trials in total, with an equal number of trials of each type. Thus, each of the four trial types - touch alone, light alone, both light and touch, and no stimulus - occurred 51 times. After the 20 ms stimulus period, there was a further 750 ms of blank screen. Finally, a response screen appeared asking the participant if they had felt a touch. Participants were asked to respond using the button box held in their right hand with one of four buttons to indicate “Definitely yes”, “Maybe yes”, “Maybe no”, or “Definitely no”. The response screen disappeared when the response was made. No feedback was provided. Finally, the screen remained blank for 1 to 1.5 s before the next trial.

### Data analysis

We first performed three analyses using a standard ANOVA framework. These analyses were performed primarily for comparison with previous studies using the SSDT, which used standard ANOVA analyses of touch reporting rates and of the signal detection measures sensitivity (*d′*) and response criterion (*c*). For all analyses, we combined “Definitely yes” and “Maybe yes” into “yes” reports and “Definitely no” and “Maybe no” into “no” reports.

For the analysis of Type-I signal detection measures, we calculated *d′* and c separately for trials with and without a light, and during active and sham stimulation. “Yes” reports on touch trials were hits; “yes” reports on no touch trials were false alarms. “No” reports on touch trials were misses; “no” reports on no touch trials were correct rejections. Thus, we had four *d′* and four *c* measures for each participant. The log-linear correction was used to account for cells with either 100% or 0% reports of touch. For the analysis of reporting rates, we ran a repeated-measures ANOVA with the factors Touch (Touch/No touch), Light (Light/No light), and Stimulation (Active/Sham) with the percentage of reports of touch as the dependent variable. Where necessary, post-hoc t-tests with Bonferroni-Holm correction for multiple comparisons were conducted to decompose significant interactions.

In addition to our standard ANOVA analyses, we also fitted a Bayesian generalized linear mixed effects model with a logistic link function using the *brms* package (see below). A key advantage of using a logistic link function is that it appropriately models the change in variance over the response scale: as mean reporting rates approach 100% or 0%, the variance decreases. ANOVA conducted on percentages does not account for such changes in variance and can lead to misleading conclusions [20]. We coded “yes” responses as 1 and “no” responses as 0, combining “Definitely yes” and “Maybe yes” into “yes” reports and “Definitely no” and “Maybe no” responses into “no” reports. The model contained three fixed effects factors - Stimulation (Active or Sham), Touch (Touch or No touch), and Light (Light or No light) - and all interactions between them. Participant was specified as a random effect, with random slopes for each fixed effect and all interactions, random intercepts, and a full, unstructured correlation matrix. The model was fitted using a No-U-Turn Sampler, a Monte-Carlo Markov-Chain (MCMC) algorithm implemented in Stan [21].

We used non-informative priors in our analysis. Specifically, there were improper uniform priors from negative to positive infinity on the mean for population-average (i.e. fixed) effects, including the intercepts; an LKJ-prior (v = 1) on the correlations between the random slopes and the intercept; and a half (i.e. constrained to be positive) student-*t* prior with shape parameter 3 and scale parameter 10 on the standard deviations of the random slopes. These priors provide little information regarding the parameter values, primarily serving to regularize the estimates of the parameters of the random effects structure. This ensures that all parameters are identifiable, and biases them against reaching improbably large values. We ran four Markov chains simultaneously, each for 5000 iterations. The first 2500 of those iterations were discarded as warm-up samples to adaptively tune the MCMC sampler. Convergence of the chains was assessed by visual inspection of their traces, which indicated that they mixed well and converged on the same parameter spaces. The *R* statistic [22] was ∼1.00 for all parameters.

In a Bayesian framework, the MCMC sampler produces a posterior distribution of likely parameter values, which we summarise using 95% credible intervals. Where necessary, we also calculated the ratio of posterior samples below zero relative to those above zero. Ratios above one indicate that more posterior samples were below zero than above it, while ratios below one indicate that more posterior samples were above than below zero. Larger values indicate high probability in favour of the hypothesis, and smaller values indicate high probability in favour of the alternative hypothesis.

All analyses were conducted using R [23] and the R-packages *afex* [24], *brms* [25,26], *emmeans* [27], *metaSDT* [28], *papaja* [29], *survival* [30], *tidybayes* [31], and *tidyverse* [32].

## Results

We first examined performance in a classical SDT framework. We found no significant difference in sensitivity (*d′*) between trials with a light (1.70) and trials without a light (1.79, [*F* (1, 20) = 2.07, *MSE* = 0.08, *p* = .165, 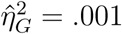]), and no significant effect of Stimulation on *d′* [Sham = 1.69; Active = 1.80; F (1, 20) = 0.16, MSE = 1.59, p = .693, 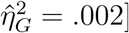]. There was also no significant interaction between Stimulation and Light on d’ [*F* (1, 20) = 1.04, *MSE* = 0.04, *p* = .319, 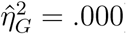], see Figure 1a.

**Figure 1.**
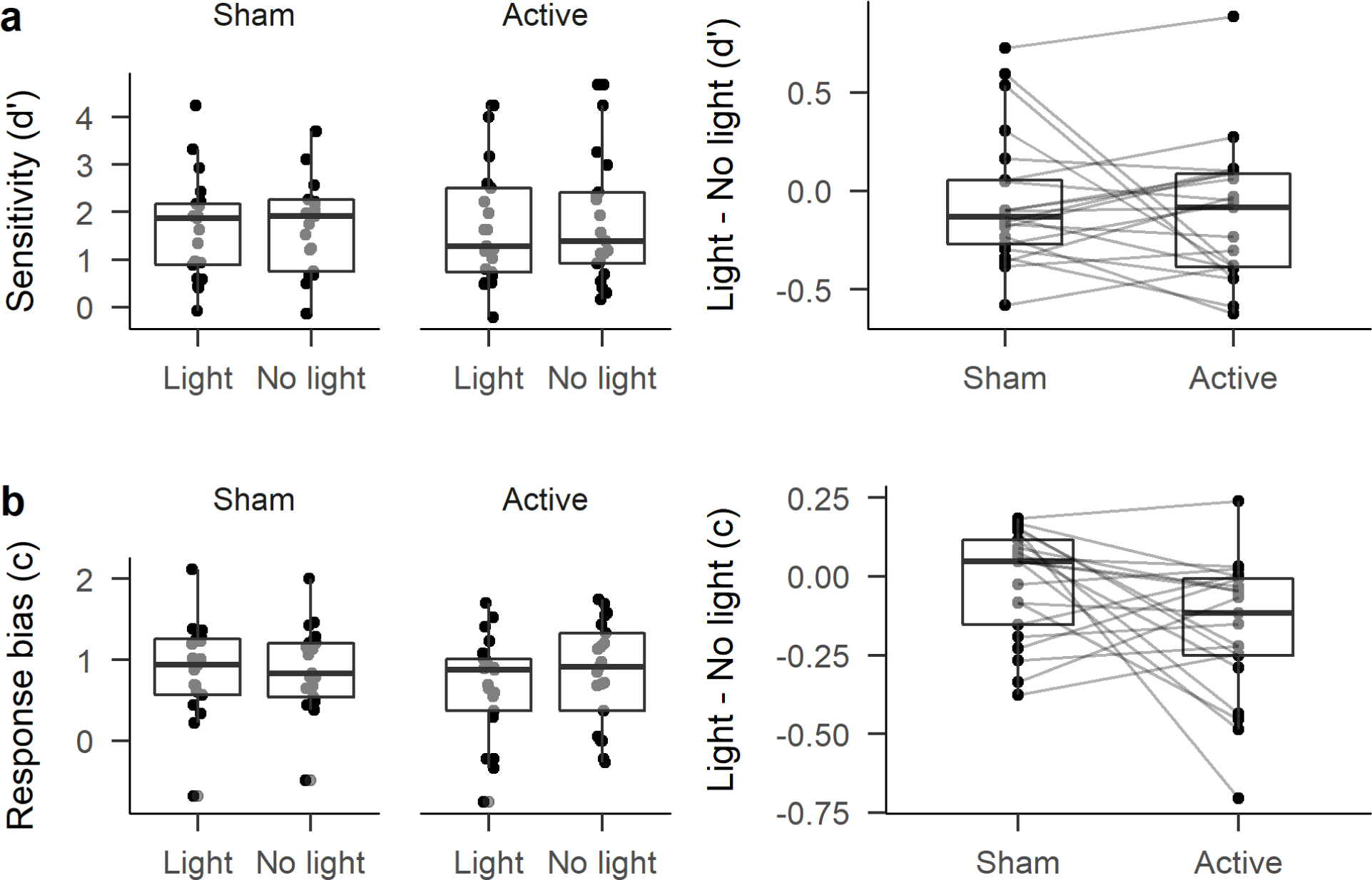
Boxplots of the signal detection measures d’ (row a) and c (row b). Boxes indicate the inter-quartile range. Lines within the boxes indicate the median. Whiskers extend 1.5 times above and below the inter-quartile range. Individual dots show individual participant scores. The right column shows the difference between d’ and c in the Light and No Light conditions in order to show the interaction between light and stimulation. Lines connecting individual dots join data points belonging to the same participant.

For response criterion (*c*), there was no significant main effect of Stimulation (Sham = 0.89; Active = 0.76; [*F* (1, 20) = 1.02, *MSE* = 0.35, *p* = .325, 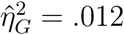]). However, there was a significant main effect of Light [*F* (1, 20) = 10.03, *MSE* = 0.02, *p* = .005, 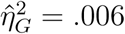], with a more liberal bias (i.e. an increase in “yes” reports) on light trials (*c* = 0.77) than on no light trials (*c* = 0.87). Importantly, there was a significant interaction between Stimulation and Light [*F* (1, 20) = 5.16, *MSE* = 0.02, *p* = .034, 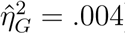], see Figure 1b. This interaction was driven by a significant difference between light and no-light trials in the Active stimulation condition (*p* = .001, Bonferroni-Holm corrected for 6 comparisons). Specifically, there was lower *c* on trials with a light (*c* = 0.67) than on trials with no light (*c* = 0.84). In Figure 1b, the pattern of lines in the interaction plot suggest a degree of heterogeneity in the interaction between Stimulation and Light for response criterion, but with the most consistent change being a shift towards a more liberal response criterion for light trials relative to no light trials (i.e. more negative values). No other comparisons were significant (all *p*s = 1).

In our analysis of reporting rates, there was a significant effect of Touch [*F* (1, 20) = 57.56, *MSE* = 0.13, *p* < .001, 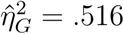], with reports of touch much more likely on trials with touches (48.51%) than without (6.26%). No other effects were significant (all *p*s > .06; see Table 1 and Figure 2).

**Table 1.**
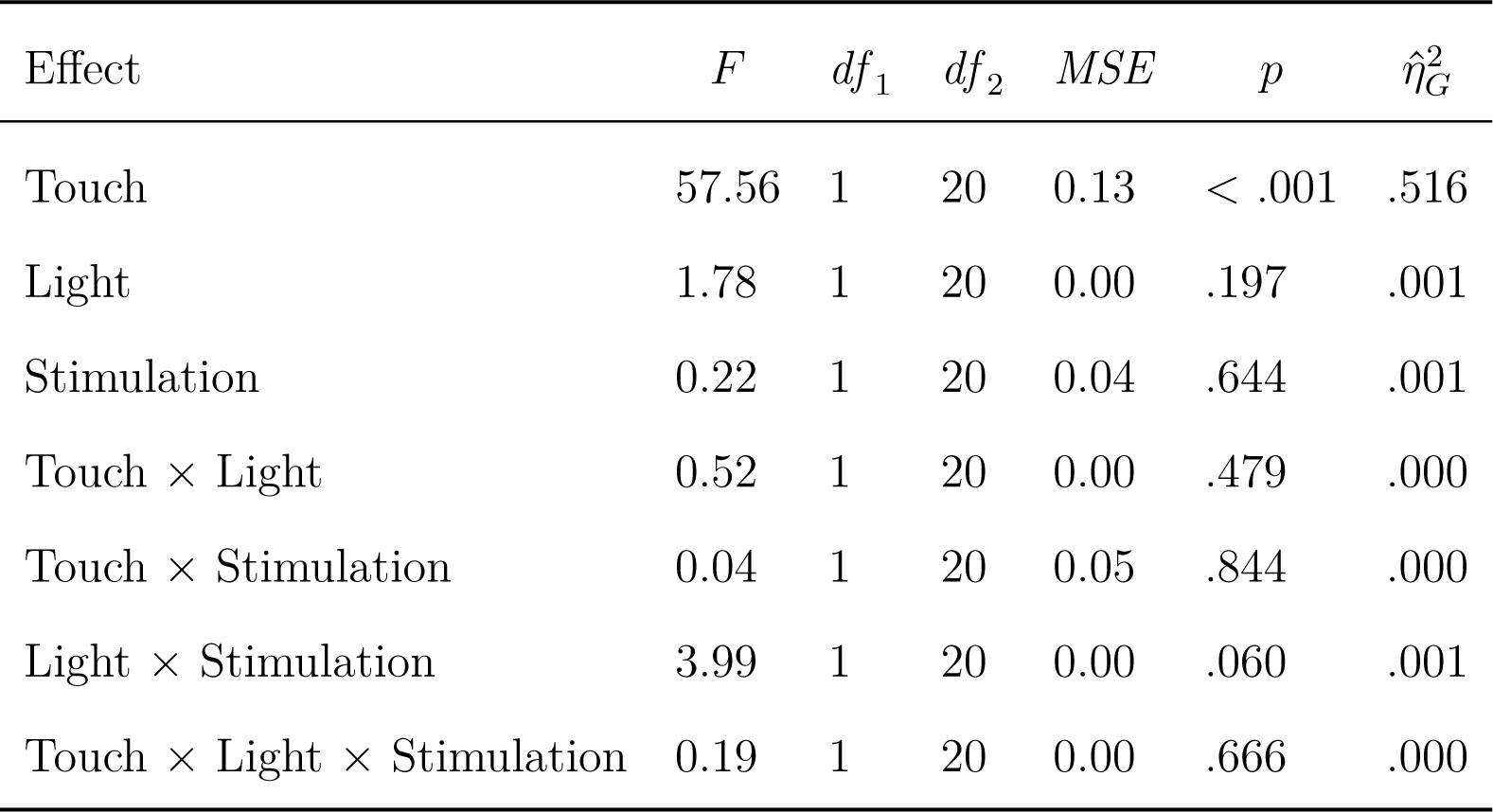
Results of the repeated measures ANOVA on touch reporting rates.

| Effect | $F$ | $df_1$ | $df_2$ | $MSE$ | $p$ | $\hat{\eta}_G^2$ |
| --- | --- | --- | --- | --- | --- | --- |
| Touch | 57.56 | 1 | 20 | 0.13 | < .001 | .516 |
| Light | 1.78 | 1 | 20 | 0.00 | .197 | .001 |
| Stimulation | 0.22 | 1 | 20 | 0.04 | .644 | .001 |
| Touch × Light | 0.52 | 1 | 20 | 0.00 | .479 | .000 |
| Touch × Stimulation | 0.04 | 1 | 20 | 0.05 | .844 | .000 |
| Light × Stimulation | 3.99 | 1 | 20 | 0.00 | .060 | .001 |
| Touch × Light × Stimulation | 0.19 | 1 | 20 | 0.00 | .666 | .000 |

**Figure 2.**
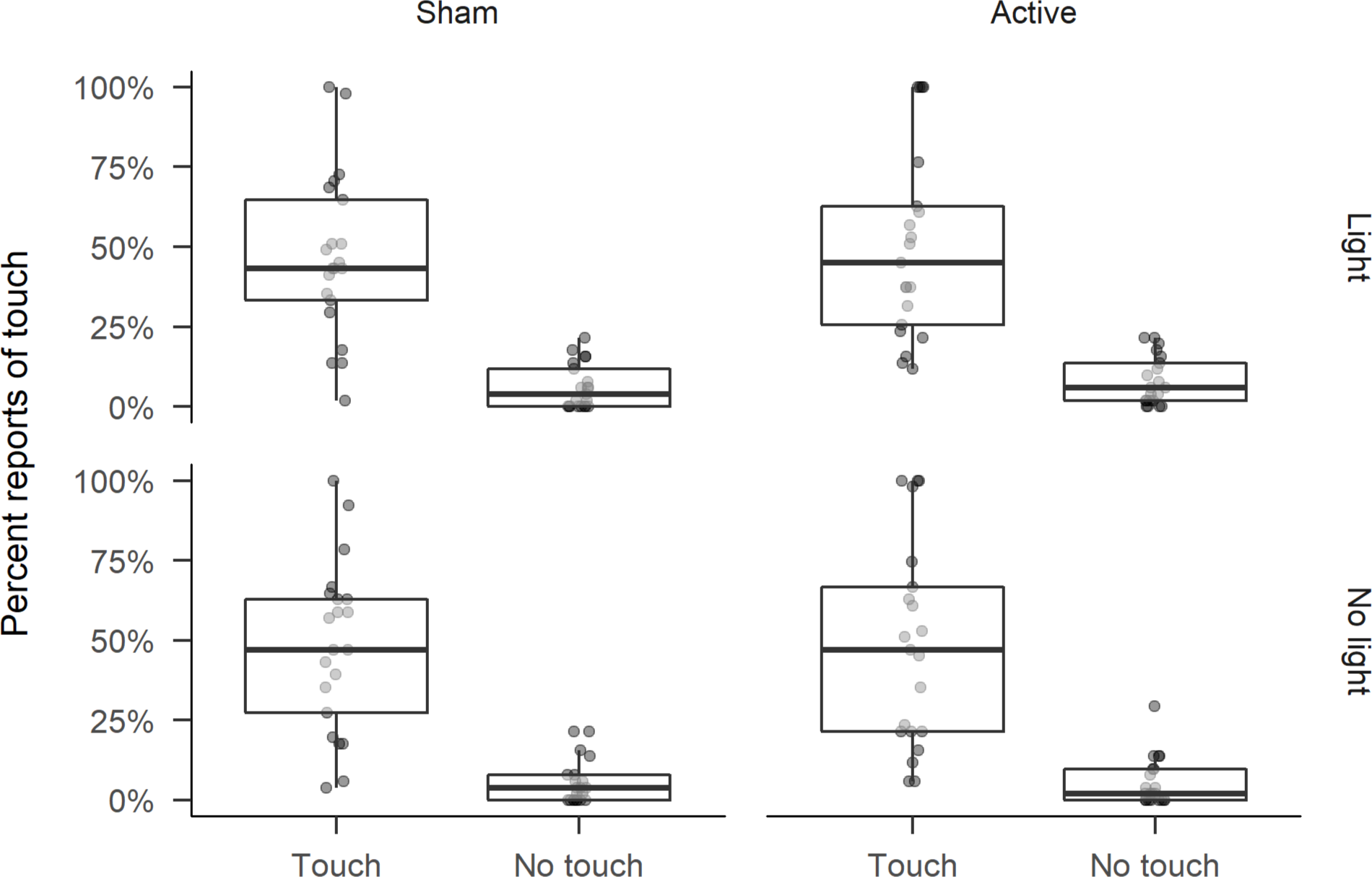
Boxplots of mean response rates in each combination of stimulation, touch and light conditions. Boxes indicate the inter-quartile range. Whiskers extend 1.5 times above and below the limits of the inter-quartile range. Lines within the boxes show the median. Individual dots indicate mean response rates for individual participants.

### Bayesian multilevel model

The Bayesian GLMM proved notably different from the repeated measures ANOVA on reporting rates (see Table 2 and Figure 3). The strong effect of Touch on reporting rates was consistent with the ANOVA, but the model also suggests that there was a small increase in reporting rates on Light trials, with the vast majority of posterior samples for this coefficient being above zero (*p*(*β* <0) = 0.02). Furthermore, the interaction between Light and Touch was also strongly likely to be negative (*p*(*β* < 0) = 21.42). On touch trials, the difference between light and no light trials was inconsistent, sometimes positive, sometimes negative. On no touch trials, reporting rates were consistently higher on light trials than on no light trials (see Figure 4a).

**Table 2.**
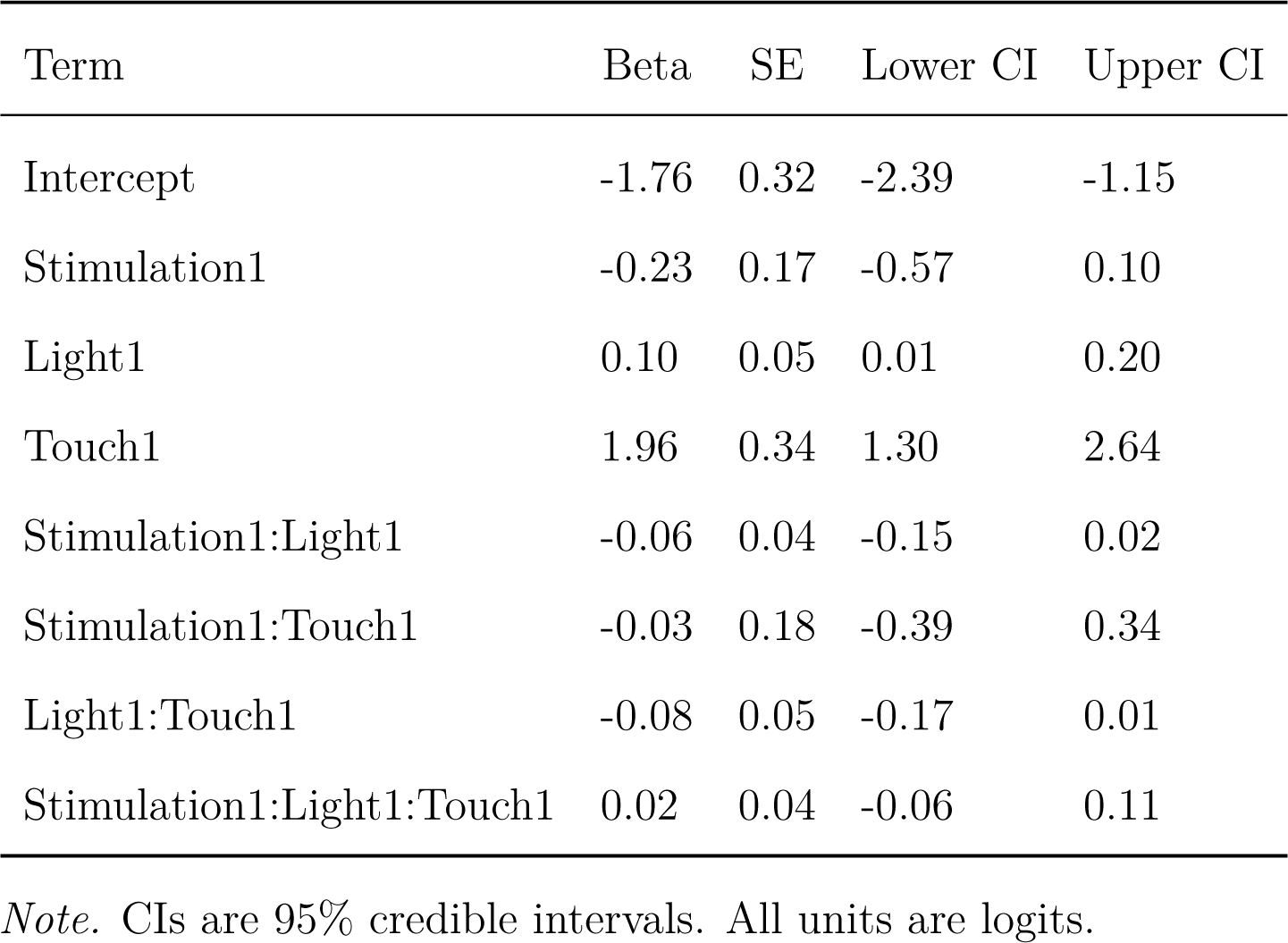
Table of fixed effects from the Bayesian GLMM.

| Term | Beta | SE | Lower CI | Upper CI |
| --- | --- | --- | --- | --- |
| Intercept | -1.76 | 0.32 | -2.39 | -1.15 |
| Stimulation1 | -0.23 | 0.17 | -0.57 | 0.10 |
| Light1 | 0.10 | 0.05 | 0.01 | 0.20 |
| Touch1 | 1.96 | 0.34 | 1.30 | 2.64 |
| Stimulation1:Light1 | -0.06 | 0.04 | -0.15 | 0.02 |
| Stimulation1:Touch1 | -0.03 | 0.18 | -0.39 | 0.34 |
| Light1:Touch1 | -0.08 | 0.05 | -0.17 | 0.01 |
| Stimulation1:Light1:Touch1 | 0.02 | 0.04 | -0.06 | 0.11 |

*Note.* CIs are 95% credible intervals. All units are logits.

**Figure 3.**
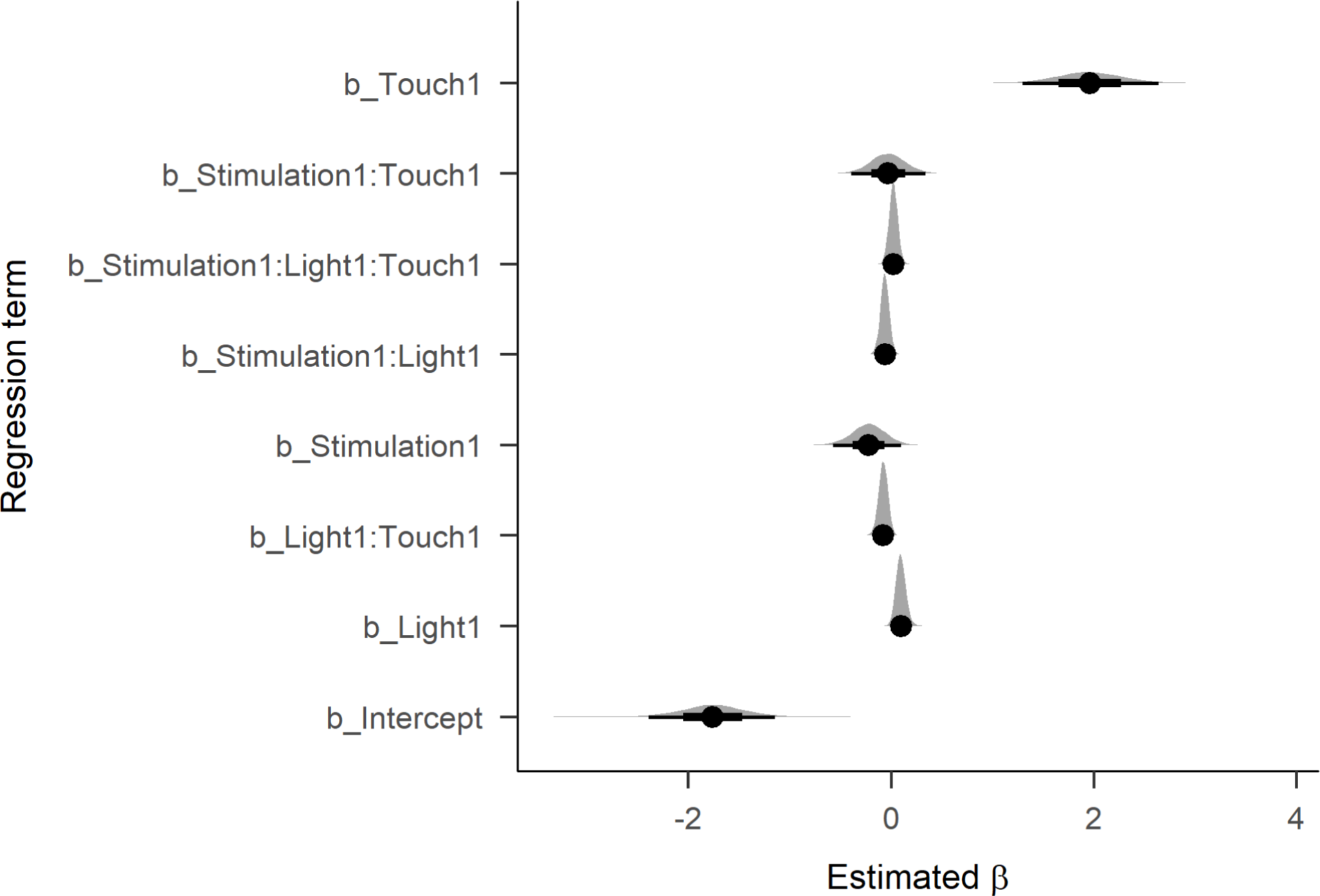
Posterior densities and credible intervals for the fixed effect coefficients. Dots indicate the mean of the posterior distribution. Bars indicate 66% (thick) and 95% (thin) credible intervals.

**Figure 4.**
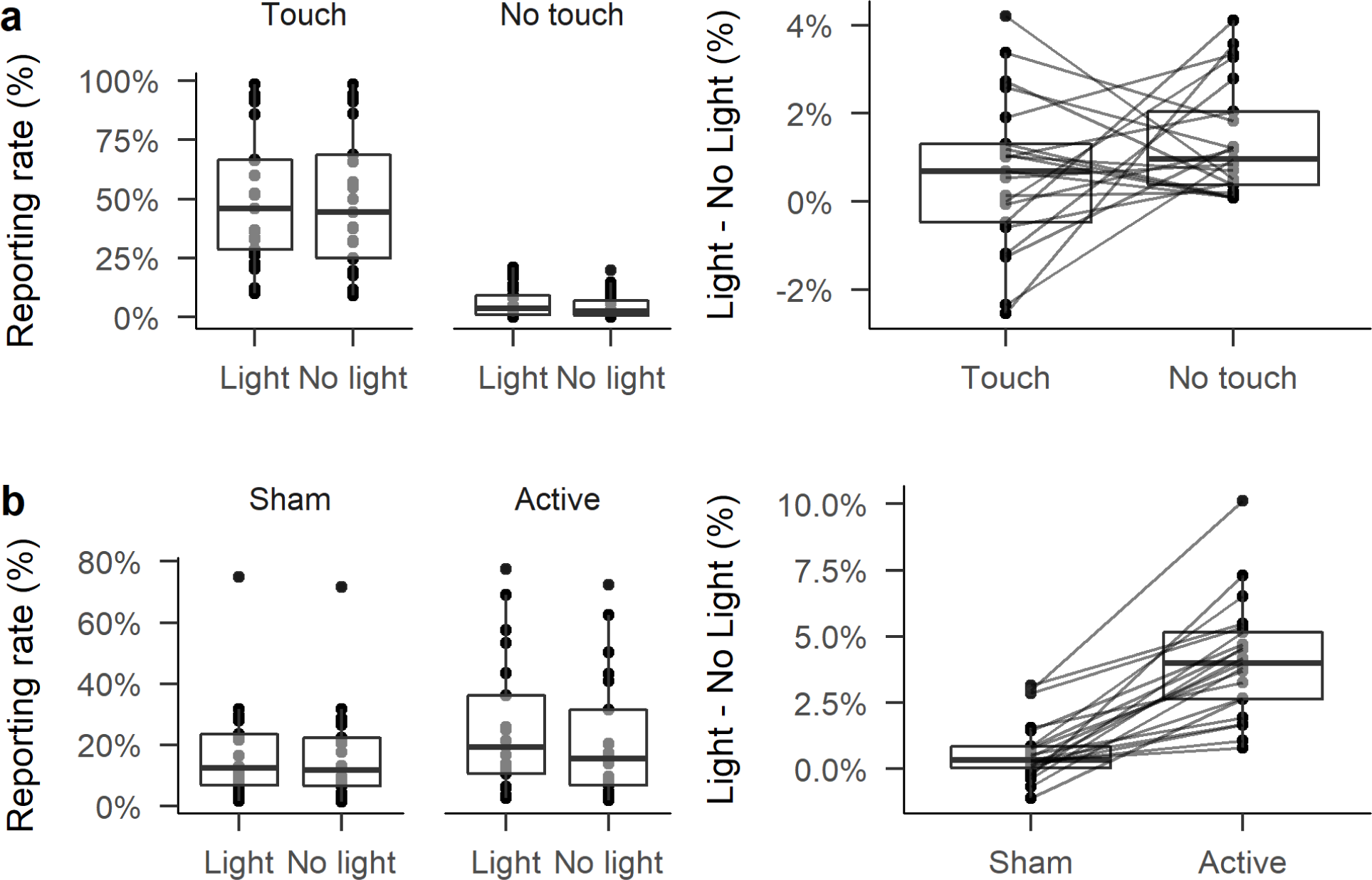
Boxplots showing model predicted yes-response rates (left) and the percentage point difference in yes-response between Light and No light trials (right). Boxplots span the interquartile range of the data, with the median shown by a single line. Whiskers extend 1.5 times the IQR above and below the hinges of the boxes. Each dot represents predicted values for individual participants. Lines join predictions from individual participants.

More importantly, the model also suggested that some Stimulation effects were also non-zero. The coefficient for the effect of Stimulation was negative (*β* = −0.23), and most of the posterior density fell below 0 (*p*(*β* < 0) = 10.70), indicating that the coefficient has a high probability of being below zero. Thus, reporting rates were likely higher overall in the Active condition than in the Sham condition. Importantly, the interaction between Stimulation and Light, though small, was also likely to be negative (*β* = −0.06, CIs = [−0.15, 0.02], *p*(*β* < 0) = 12.14). As can be seen in Figure 4b, on Sham stimulation trials, there were was little difference between trials with a light and without a light. But during Active stimulation, all participants showed increased reporting of touches during trials with a light compared to trials without a light.

Critically, there was little evidence of an interaction between Stimulation and Touch. The posterior density spanned zero, with only a low probability of the parameter being negative (*p*(*β*) < 0 = 1.32). The three-way interaction between Stimulation, Touch, and Light was similarly equivocal, with a posterior ratio more in favour of the parameter being positive than negative (*p*(*β*) < 0 = 0.41). Thus, to the extent that Stimulation had effects on reporting of touch, these effects were driven by changes in responses to the light.

## Discussion

We examined the effects of 10 Hz transcranial alternating current stimulation (tACS) over centro-parietal regions on performance of the Somatic Signal Detection Task. We previously reported that oscillatory activity in this frequency range influenced reporting of touch independently of whether touch is actually present [3]. Our analysis of signal detection measures suggested that tACS stimulation did not influence detection sensitivity, but did introduce a more liberal bias towards responding that touch was present, particularly in the presence of flashes of light. Our Bayesian model also suggests that reports of touch were increased during active stimulation, with an additional increase in the presence of light flashes. This was independent of whether a target touch stimulus was present or not. In combination, these results suggest that tACS stimulation at 10 Hz modulated response bias independently of sensitivity. As reported in Craddock et al.[3], reports of touch decline as alpha power increases and increase as alpha power decreases, independent of whether touch is present. Although 10 Hz tACS stimulation of visual cortex increases occipital alpha power, Gundlach et al.[19] reported that 10 Hz tACS stimulation decreased somatosensory alpha rhythms. A decrease in somatosensory alpha power would thus lead to an increase in reporting of touch and a more liberal response bias, which is what we found here.

An explanation for the influence of alpha power on touch is that it may reflect variation in cortical excitability [33–36]. Alpha power increases as cortical inhibition increases, and decreases with increased cortical excitability [37]. The balance of excitation and inhibition across cortical areas may reflect suppression of sensory responses during selective attention [38]. For example, during visual spatial attention tasks, oscillatory power in the alpha band is lower over the hemisphere contralateral to the attended region of space and higher over the hemisphere ipsilateral to the ignored region of space [39]. Increasing inhibition suppresses low-level cortical responses and restricts outflow of information to higher-level cortical areas [40]. Concomitantly, an increase in excitability lifts that gate and allows more information out, therefore shifting to a more liberal response bias.

Nevertheless, in the context of an increase in cortical excitability in somatosensory cortex, the interaction with the light is unexpected. We might instead have expected overall response rates to increase irrespective of the influence of the light. However, the effect of light was multiplicative with active stimulation. Active stimulation increased reports of touch even without the light; the increase was simply larger when the two were combined. During sham stimulation, there was little consistent difference in reporting rates between light and no-light trials. Thus, the combination of both active stimulation and light flashes induced a more liberal response bias. An increase in output from somatosensory regions would give increased opportunities for the light to boost responses to perceived somatosensory stimulation.

Our results do come with some caveats. First, our comparison of active versus sham stimulation would not allow us to make concrete statements about the specificity of stimulation at a particular frequency, since we stimulated only at a single frequency. Second, since we did not record EEG before and after stimulation, we cannot be sure that we directly influenced visual alpha or somatosensory alpha rhythms. Finally, since we used only a single pair of stimulation locations, we cannot necessarily distinguish between non-specific effects of tACS stimulation and direct effects of stimulation on the specific rhythms of interest. Overall, however, our results are consistent with tACS stimulation at 10 Hz over somatosensory regions altering response bias in the SSDT, and thus provide support for a direct role of alpha oscillatory rhythms in tactile perception.

## Acknowledgements

The authors thank Jumanah Hendrickson-Wilson and Hannah Clark for assistance in collection of the data.

*Funding:* This study was funded by the Biotechnology and Biological Sciences Research Council (BBSRC; BB/L006618/1). Wael el-Deredy received funding from CONICYT, Chile, Basal project.

